# Variations in microbial diversity affect the stability and function of dark fermentation bioreactors

**DOI:** 10.1101/2022.06.27.497814

**Authors:** Marcelo Navarro-Díaz, Valeria Aparicio-Trejo, Idania Valdez-Vazquez, Julián Carrillo-Reyes, Morena Avitia, Ana E. Escalante

## Abstract

The relationship between the taxonomic diversity and the function of microbial communities is complex. Specifically, the ecological mechanisms that drive the dynamics of microbial populations and the consequences of these dynamics on functional traits have remained elusive. Among the simplest but natural microbial communities are dark fermentation consortia, a subset of the more diverse and complex microbial communities, anaerobic digestion communities. Dark fermentation consortia have been of interest as they produce biofuels such as hydrogen and different alcohols that can be used as fossil fuels alternatives. However, these hydrogen-producing communities have unresolved instability and low yield issues. We have previously proposed that instability and low yields in dark fermentation communities could be due to reduced diversity that results from aggressive pretreatments of original anaerobic digestion communities. In this work, we used dark fermentation communities to examine experimentally the effect of diversity reduction in functional traits, including stability and microbial interactions. We established two types of treatment, (i) maintaining strict culture conditions that are known to induce hydrogen production and ii) applying a heat-shock treatment known for selecting hydrogen-producing bacteria, which resulted in two types of communities, high and low diversity. Each treatment consisted of 12 replicates that were transferred to fresh medium daily (during 28 days for the non-treated bioreactors and 61 days for the heat-shock treated bioreactors). Microbial communities of the two treatments were characterized in their function as well as resistance to invasion. Microbial composition was characterized by culture-independent 16S rRNA gene amplicon sequencing. We analyzed microbial community composition and function through time, establishing statistical relationships between bacterial groups and metabolite production. Also, we inferred the potential ecological interactions that might have been established. Results show that the replicate bioreactors for each treatment predictably shifted to a similar composition and increased and stable biogas production. The non-treated bioreactors showed less susceptibility to the invasion (with the invasive bacteria establishing only in one replicate in the non-treated bioreactors vs 6 invaded replicates of the heat-shock treated bioreactors). However, the effect observed in the non-treated bioreactor replicate where the invader bacteria established was more drastic since the invasive bacteria managed to become dominant.

## Introduction

The high taxonomic and metabolic diversity of bacterial groups give rise to complex functional behaviors in microbial communities where even thousands of different bacteria might coexist (Locey and Lennon, 2016; Louca et al., 2018). The relationship between the microbial composition, diversity and their functional outcomes can be difficult to predict and control since these result from complex metabolic and environmental interactions (Escalante et al., 2015; Hays et al., 2015). Also, microbial communities are susceptible to perturbations (i.e. climate change, pollution and nutrient availability) which might change their composition and functioning (Allison and Martiny, 2009). Moreover, the mechanisms behind such changes have been observed to be case-specific (Shade, 2017). It has been suggested that the increase in diversity components (i.e. richness and evenness) leads to higher stability and functional resilience due to functional redundancy, niche complementation and differential response traits against perturbations, but mixed evidence has been found (Shade et al., 2012). In bioreactors, such as those from dark fermentation and anaerobic digestion, it has been found that consortia with higher alfa-diversity were more resilient to environmental pH disturbances (Feng et al., 2017), while evenness has been found to increase stability by allowing the community more capacity to use a varied array of metabolic pathways (Werner et al., 2011). Similarly, in soil bacterial communities increased diversity was related to increased stability against heat perturbations (Tardy et al., 2014; Xu et al., 2021). In contrast, Wertz et al. found no effect on stability against heat disturbances of decreasing bacterial diversity in bacterial communities of soil (Wertz et al., 2007) and Glasl et al. found that in marine sponges-associated bacterial communities varying levels of microbial diversity did not affect the stability against salinity disturbances (Glasl et al., 2018). Further, not only the identity and abundance of species are important for the stability and function of bacterial communities. For example, biotic interactions are important to determine the assembly and composition of bacterial communities (Pérez-Gutiérrez et al., 2013; Datta et al., 2016; Friedman et al., 2017; Meroz et al., 2021). Besides, these interactions are important in determining the function, stability and invasibility of communities (Ghosh et al., 2016; Kinnunen et al., 2016; Madsen et al., 2018; Ratzke et al., 2020). Since microbial communities carry out important processes like nutrient cycling, substrate degradation and metabolites production, understanding the mechanisms that drive the diversity-function relationship can help in the management of microbial communities, for example, in biotechnological o bioremediation settings (Johns et al., 2016).

Using microbial consortia with reduced complexity can be the first step to understanding the mechanisms that link diversity and function. Simplified microbial consortia (like those used for biofuels production or artificially-assembled consortia) have been used to test ecological hypotheses and propose generalizable ecological theories since they offer controlled environments and easily measurable functions (De Roy et al., 2014; Stenuit and Agathos, 2015; Cairns et al., 2018). Due to the intrinsic characteristics of microbial organisms (i.e. their small size and chemically-mediated interactions; Schmidt et al., 2015) the exploration of ecological mechanisms (e.g. biotic interactions) in microbial communities is frequently performed via statistical inference. In this line, to advance the knowledge regarding the precise ecological dynamics of microbial consortia, specifically designed experiments and systematized research can be performed (Navarro-Díaz et al., 2020). Experimental controls, contrasting experimental settings and replication are useful to determine the impact of culture conditions, statistical detection of patterns and robust interpretation of results. In turn, results obtained in this manner can narrow the scope of hypotheses directing further experiments.

Hydrogen-producing microbial consortia represent a good system to study ecological hypotheses since they are low-diverse systems with well-studied measurable functions (Wang et al., 2020). Hydrogen-producing (also known as dark fermentation consortia) are microbial consortia derived from anaerobic digestion communities in which hydrogen-producing bacteria are selected by using aggressive pretreatments and strictly controlling culture conditions (Wang and Yin, 2017). In hydrogen-producing reactors, instability and low yield are still unresolved issues and proposed causes underlie ecological processes like biodiversity loss and competitive interactions (Castelló et al., 2020). It has been known that certain species present in hydrogen-producing consortia have a certain impact on the community function. For instance, *Clostridium* species have been attributed with hydrogen production and degradation of complex substrates and *Bacillus* species with eliminating toxic oxygen (Cabrol et al., 2017). Lactic acid bacteria (LAB), like *Lactobacillus* species, are of special interest in bioreactor research. Initially depicted as inhibitors of hydrogen producers by the production of bacteriocins and substrate competition, recently, they have been regarded as aiding hydrogen production and participants in detoxification processes (Sikora et al., 2013; Muñoz-Páez et al., 2018). The specifics of the roles LAB play when they colonize bioreactors are still important to elucidate, especially as we advance in the scaling process of dark fermentation where non-sterile substrates are to be used. Importantly, most of these functions have not been studied in an ecological context that can provide insights into population dynamics and community-level properties.

In this work, we studied the effect that changes in diversity had on the function and the stability of microbial consortia. For this, we used hydrogen-producing consortia as a model system. Carrying out two commonly used strategies for achieving hydrogen production, we established two sets of microbial consortia with two levels of diversity from the same inoculum. The first strategy consisted of applying an aggressive heat-shock pretreatment to the inoculum while the second consisted of maintaining specific culture conditions previously reported to allow for hydrogen production. We hypothesize that since diversity determines the function and stability of microbial consortia, each of the two sets of hydrogen-producing bioreactors will not only have varying degrees of diversity but also their long-term behavior and resistance to perturbations.

## Methods

### Experimental design

To investigate if microbial diversity affects the ecological robustness of dark fermentation microbial consortia, we used lab-scale bioreactors inoculated with anaerobic digestion granules and applied two different treatments to control for levels of diversity that permitted to evaluate the different functional outcomes. The experiment was conducted in two sequential stages, in the first stage we intended to specifically evaluate the effect of diversity on the functional stability of the bioreactors; in the second stage, we challenged the bioreactors with a controlled invasion of *Lactobacillus plantarum* culture. For the first stage, we artificially reduced the diversity of the original inoculum with a heat-shock treatment (Valdez-Vazquez and Poggi-Varaldo, 2009) to further compare the microbial community dynamics and performance of the heat-shock treated microbial communities against the non-treated communities. Twelve replicates per treatment (heat-shock treated and non-treated) were maintained with daily transfers into fresh medium and until biogas production stability was achieved. After biogas production stabilized, for each of the two “diversity treatments” we chose six random replicates to start the second stage of the experiment. The six chosen replicates per diversity treatment were divided into two new set of bioreactors. One set was inoculated with a strain of *L. plantarum* that was isolated from a hydrogen-producing bioreactor anaerobic bioreactor (Pérez-Rangel et al., 2021), the other six were used as non-invasion controls (Figure 1). The functioning or performance of the bioreactors throughout the experiments was determined based on three parameters: daily biogas production, volatile fatty acids (VFAs) concentration and pH. Microbial diversity and composition were determined by culture-independent 16S rDNA amplicon high throughput sequencing.

**Figure 1.**
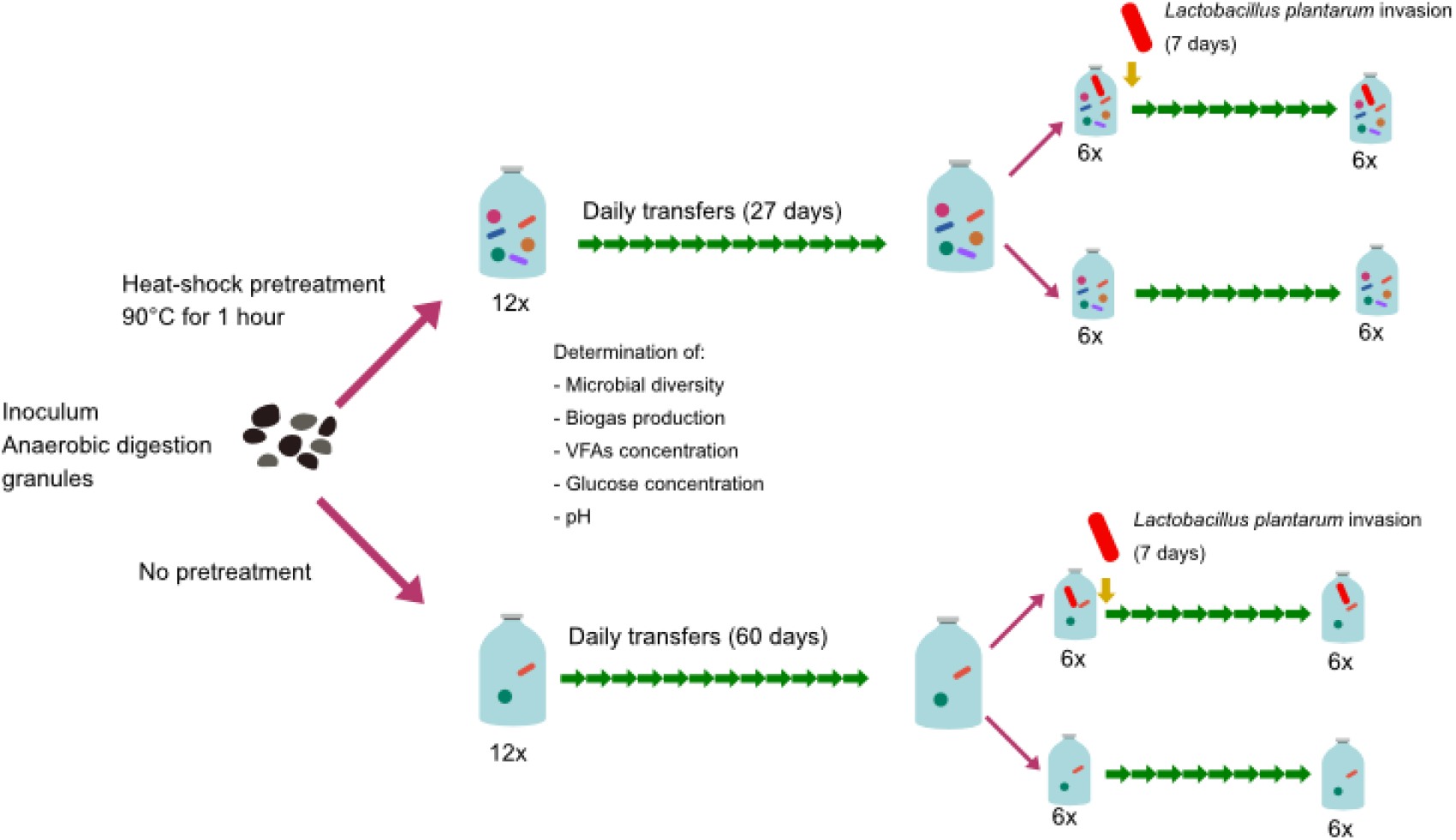
Experimental design. Starting from a single inoculum, we followed two experimental treatments to artificially modify microbial diversity. Each treatment consisted of 12 replicates derived from a single inoculum. To test the stability of reactors in each treatment, we periodically measured the performance of bioreactors until biogas production stabilized. After biogas production stabilization, we challenged the reactors with a controlled invasion and determined the success of the invasive bacteria based on their abundance in the reactors. In the invasion experiment, 6 bioreactors per treatment were selected and divided into two sets of bioreactors. One set was inoculated with *Lactobacillus plantarum* simulating a biological invasion, while the remaining 6 bioreactors per treatment were used as controls. After biogas production in invaded bioreactors stabilized, the experiment was terminated. All the data on the diversity and performance of the bioreactors were subject to statistical analyses for formal comparisons of the experimental treatments.

### Bioreactors’ setup

To compare the effect of diversity on the function and stability of bioreactors, we artificially reduced the diversity of the original inoculum with a heat-shock treatment. The original microbial inoculum consisted of anaerobic digestion granules obtained from a brewery wastewater treatment plant. Before the experiment, 15g of the inoculum was homogenized in 20mL of 1:1 PBS/glycerol solution and frozen at –80°C until use. From this inoculum, the two diversity treatments bioreactors were established as overnight cultures using 20 mL of homogenized inoculum and 40 mL of fresh medium at 37°C. The heat-shock pretreatment consisted of heating the overnight culture to 90°C for 1h. Since, after the heat-shock pretreatment, no growth was achieved under aerobic conditions, strictly anaerobic conditions had to be used in this treatment by adding 0.5 g/L of cysteine to the culture medium. Once initial cultures were established for both treatments (non-treated and heat-shock treated bioreactors), they were divided into 12 replicates per treatment. To partition each reactor, every 24hours, 20 mL of the culture were transferred into a new serum bottle with 40 mL of fresh medium. Then, 20mL of the culture in original bioreactor was kept and 40mL of fresh medium were added. This partition step was repeated 4 times (for 4 days) and 12 random replicates were used for each treatment. From that point, every 24 hours, 20mL of the culture of each replicate culture was reinoculated into 40mL of fresh medium. After each transfer, pH was adjusted to 6. The medium had the following composition per liter: glucose (5 g/L), urea (0.65 g/L), K_2_HPO_4_ (0.25 g/L), MgCl_2_·6H_2_O (0.376 g/L), FeSO_4_· 7H_2_O (0.1 g/L), CoCl_2_·6H_2_O (0.0025 g/L), MnCl_2_·4H_2_O (0.0025 g/L), KI (0.0025 g/L), NiCl_2_·6H_2_O (0.0005 g/L), ZnCl_2_ (0.0005 mg/L) and yeast extract (0.5 g/L); modified from (Mizuno et al., 2000). For all the cultures we used 100 mL serum bottles as lab-scale semi-continuous bioreactors with a working volume of 60mL. Bioreactors were incubated at 37°C and shaken at 50 rpm.

### Characterization of biogas production

To characterize the performance of the bioreactors, we measured daily biogas production. Biogas production was measured in an inverted graduated cylinder using the water displacement technique and, to ensure an exact measurement, we prevented CO_2_ absorption by using a 5N HCL solution (pH < 2; (Boshagh and Rostami, 2020). We maintained the bioreactors for at least 4 weeks or until stability was achieved. We calculated the hydrogen production stability index (HPSI; (Tenca et al., 2011)) for each replicate using the biogas production of the previous 3 days. When the HPSI mean of the 12 replicates was above 0.9 for 3 consecutive days we considered stability was achieved. We also calculated the final HPSI for each replicate and the mean of all replicates for each treatment.

### Invasion test

After the initial stabilization period, 6 random replicates for each treatment was divided again and one of the derived bioreactors for each replicate was inoculated with a strain of *Lactobacillus plantarum*. The *L. plantarum* strain was isolated from a hydrogen-producing bioreactor degrading wheat straw, operating at 37°C and pH 6.5 (Pérez-Rangel et al., 2021). The strain of *L. plantarum* was grown overnight in LB medium (Sigma Aldrich, USA) and then acclimatized for 3 days in the same medium and conditions that were used in our bioreactors. The invasion was performed at a 10% ratio. For this, we followed the same protocol for daily transfers, but instead of just 40mL of fresh medium, we used 34 mL of fresh medium and 6 mL of the overnight culture of *L. plantarum*.

### Characterization of complementary functions of bioreactors

In addition to biogas production, we measured tvolatile fatty acids (VFAs) concentration and pH as proxy measures of the function of bioreactors. Volatile fatty acids (VFAS) concentration was determined using an SRI 8610-00 gas chromatograph equipped with a Porapak Q column and using Helium at a 30mL/min flow rate as carrying gas. Temperatures for the injector, column and oven were 150C°, 50C° and 50C° respectively.

### Characterization of microbial diversity

At the same time points in which chemical analyses were performed, we obtained samples for microbial composition analyses. Samples were stored at -80°C until processing. DNA was extracted using the Quick-DNA Fungal/Bacterial DNA Microprep Kit (Zymo Research, USA) following the manufacturer’s instructions. Sequencing library preparation was performed targeting the V4 region of the 16S SSU rRNA gene using primers 515F: GTGYCAGCMGCCGCGGTAA (Parada et al., 2016) and 806R: GGACTACNVGGGTWTCTAAT (Apprill et al., 2015). Amplifications were performed in 25 µL reactions with the Platinum Hot Start PCR Master Mix (Invitrogen, USA) 0.5 µL of 10 mM primers, and 1 µL of DNA template. Negative controls were included for each PCR reaction to ensure that no contamination occurred. Later, amplification products were visualized using electrophoresis gels. Finally, the sequencing reaction was performed using the Miseq 300PE platform at the Macrogen facilities (Korea) using Reagent V3 and Phix at 30% and with a red length of 250PE.

### 16S rRNA sequence data processing

#### De-multiplexing, filtering, and chimera check

Raw sequences (10,540,264) were demultiplexed using the *demuxbyname* script from the BBTools suite (Bushnell B., sourceforge.net/projects/bbmap/). Demultiplexed sequences were next processed using QIIME 2 v2021.2 (Bolyen et al., 2019). Sequence quality control (denoising and chimera check) and inference of amplicon sequence variants (ASVs) was performed using the *dada2 denoise-paired* command. We modified the *trim left f, trun len f, trim left r*, and *trun len r* parameters to remove 30bp from both extremes of the reverse and forward sequences. After quality control, 7,893,591 (74.89%) sequences were retained. The raw data (paired-end files) were deposited in the NCBI sequence read archive (SRA) with the accession number: PRJNA814689.

#### ASV assignment

After quality filtering, samples were rarefied to the average of reads per sample (27,082 reads) using the *qiime feature-table rarefy* command with the *--p-with-replacement* argument in QIIME 2 v 2021.2. Taxonomy was assigned to 115 ASVs using BLAST (Altschul et al., 1990) against the NCBI’s 16S ribosomal RNA refseq database (O’Leary et al., 2016). Finally, to improve pattern recognition and reduce technical variability, ASVs with less than 20 reads in 20% of the samples were filtered out (retaining 24 ASVs) of the following analyses based on the filtering criteria published previously (filtering ASVs with less than *m* = 20 counts in at least *k* = 20 samples, where m and k were selected as 0.1% of the minimum sample library size which was ∼20,000; Cao et al., 2021).

### Statistical analyses

To assess the variability of microbial function over time, for each treatment, we plotted the 12 bioreactors function data (biogas production and AGVs). Similarly, to assess the variation of the average microbial abundance over time we used stacked bar plots of the average abundance of each ASV for the 12 replicates. To investigate how low-abundance ASVs behaved over time the average abundance was plotted in a heatmap, normalizing ASVs abundance across samples. To analyze the relationship between microbial composition and bioreactors we followed two approaches. First, we performed Spearman correlations between ASVs abundance counts and metabolites concentrations and biogas production obtained using the *cor* function in the *stats* R package v4.0.5. We represented the correlations as heatmaps for visual interpretation. Then, we used the *cca* function in the vegan R package v.2.5-7 (Oksanen et al., 2013) to perform a canonical correspondence analysis (CCA) using the abundance counts of ASVs and normalized metabolites concentrations and biogas production. All analyses were performed using R v.4.0.5 (R Core Team, 2016). Finally, to infer the potential interactions occurring between the members of the hydrogen-producing consortia, we used the Lotka-Volterra-based network inference approach implemented in MetaMIS v1.02 (Shaw et al., 2016). For each sampling time point, the average of the non-normalized microbial abundance of the 12 replicates was computed and used to infer ecological networks. The networks were visualized using Cytoscape 3.0 (Shannon et al., 2003).

## Results

### Bioreactor’s function

For both treatments (the treated and the non-treated treatments), hydrogen production showed increased production and stability over time being the main difference between treatments that the heat-shock treated bioreactors showed a higher biogas production overall (Figure 2). For the non-treated bioreactors, initially, hydrogen production was approximately 250 mL/L, and at the end of the experiment, it reached 800 mL/L. Also, biogas production stabilized towards the end of the experiment, such that all replicates showed similar productivity both on contiguous days and between replicates. For the treated bioreactors, high and stable biogas production was quickly achieved and showed a smaller increase since the 12 replicates produced around 1500 mL/L of biogas stabilizing at the end of the experiment around 1700mL/L. Importantly, HPSI was higher in the treated than in the non-treated bioreactors (0.904 vs 0.614). In both treatments, the concentration of VFAs remained mostly stable over time (Figure 3). The prevalence of acetic and butyric acids indicates that hydrogen production was occurring although gas composition was not measured. Overall, all replicates from both treatments stabilized in terms of biogas production several weeks after the start of the experiment and reached different production peaks depending on the treatment. In the non-treated bioreactors, concentration of acetic and butyric acids increased initially and stabilized towards the end of the experiment. Contrarily, in the treated bioreactors, VFAs and biogas showed the same pattern of stable production.

**Figure 2.**
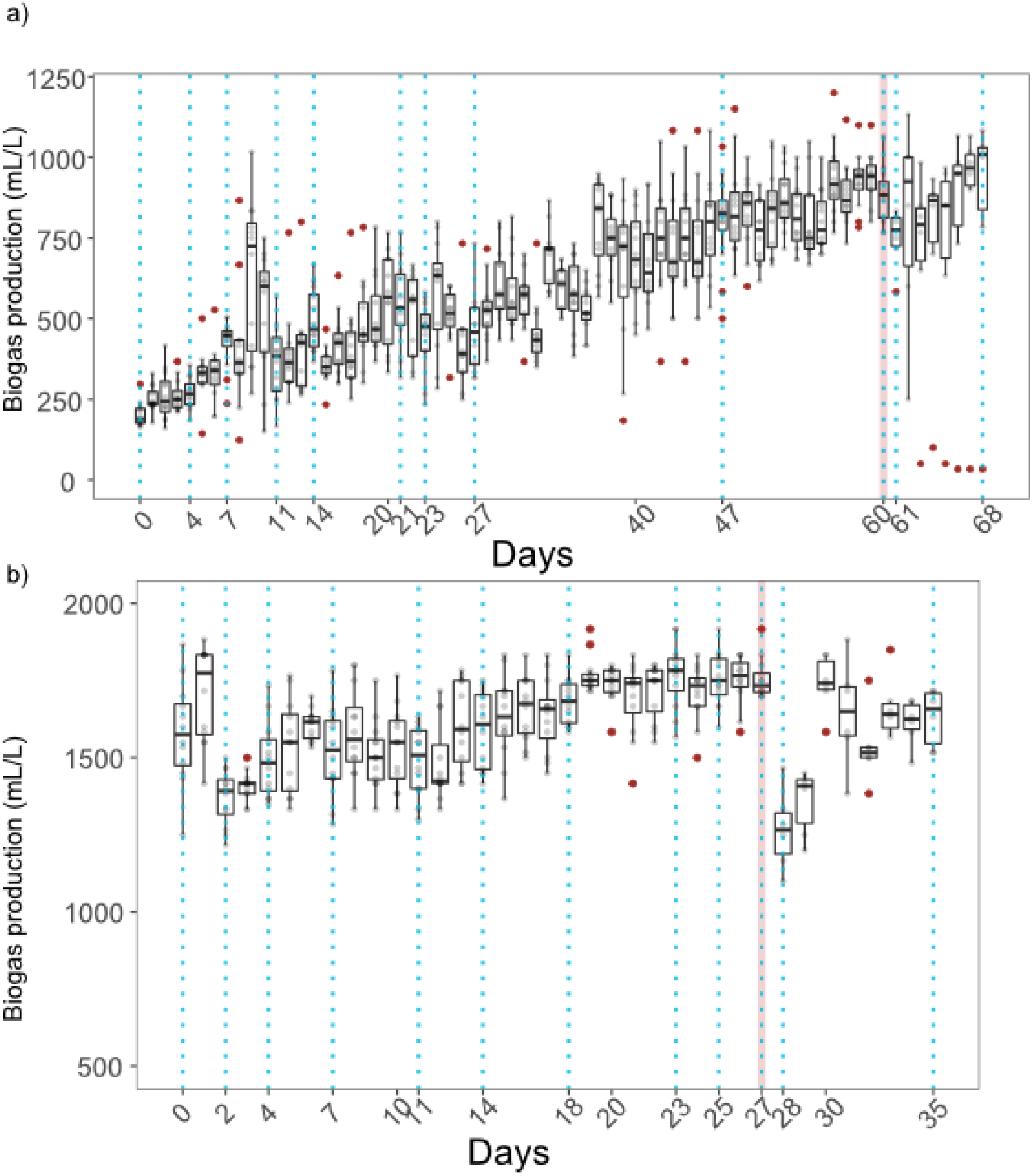
Biogas production over time in the non-treated (panel a) and heat-shock treated (panel b) bioreactors. Dotted vertical lines represent the sampling days for microbial and VFAs characterization. The red vertical line represents the day where the invasion with *L. plantarum* occurred. 12 replicates were measured per day until the invasion day where only the 6 invaded samples are shown.

**Figure 3.**
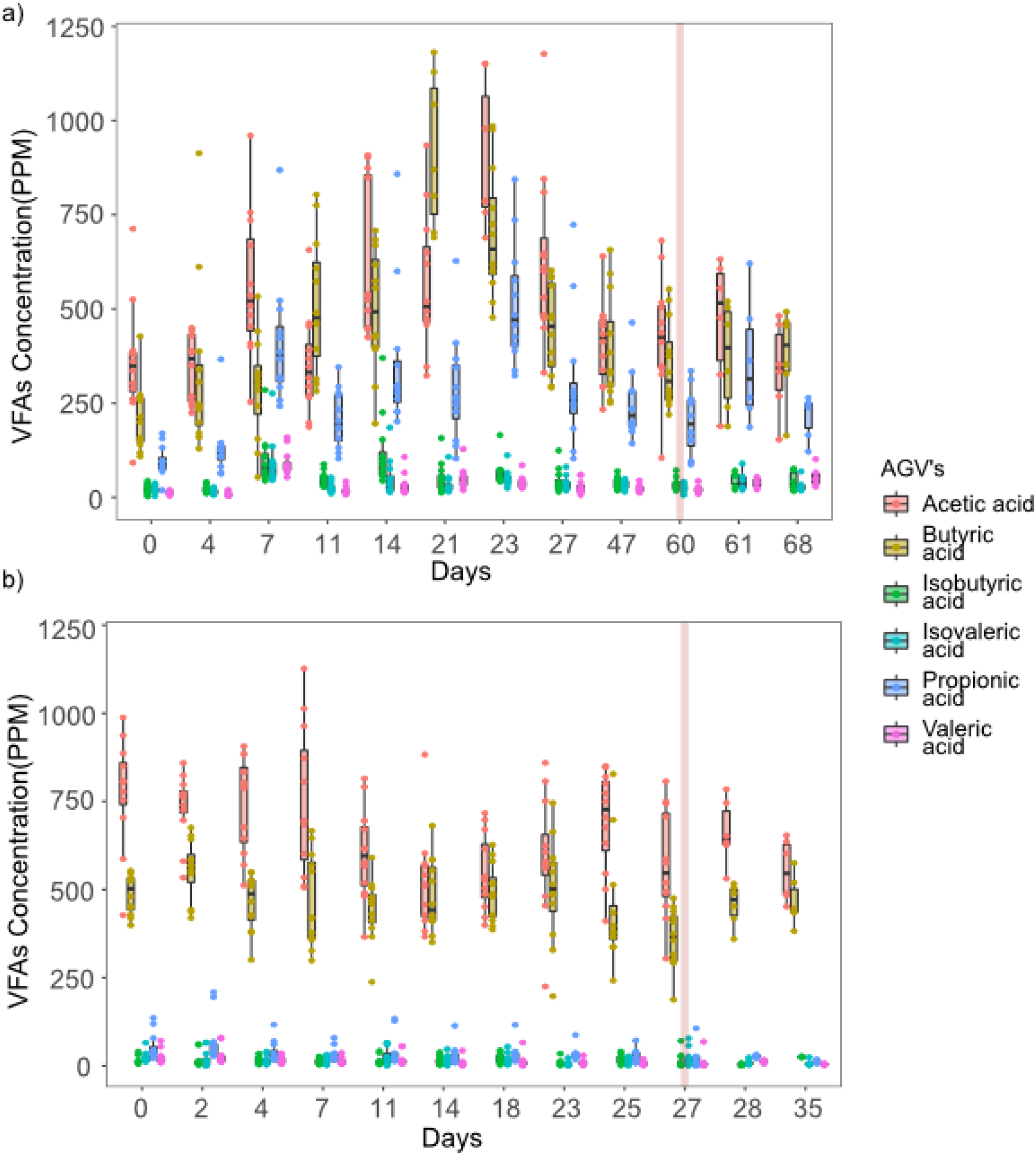
Volatile fatty acids production over time in the non-treated (panel a) and heat-shock treated (panel b) bioreactors. The red vertical line represents the day where the invasion with *L. plantarum* occurred. 12 replicates were measured per day until the invasion day where only the 6 invaded samples are shown.

The effect of the invasion by *L. plantarum* on the function of the bioreactors was different in each of the treatments. In both experiments, invasion induced a reduction in biogas production observed 24 hours after the inoculation (Figure 2). The reduction was steeper in the heat-shock treated bioreactors than in the non-treated bioreactors (72.15% vs 85.03% of the original biogas production). After the initial reduction, biogas production increased again to the similar levels observed before the invasion (93.54% and 92.87% of the original biogas production in the non-teated vs the heat-shock treated bioreactors respectively). However, in the non-treated bioreactors, one replicate did not recover from invasion and stopped producing biogas. Also, for the heat-treated bioreactors, biogas production recovery was faster than in the non-treated consortia. The concentration of the analyzed VFAs was not seemingly affected by the invasion in either treatment (Figure 3).

### Microbial composition

In the non-treated bioreactors, the microbial consortia were composed of several bacterial ASVs that showed a distinctive pattern of change during the experiment. Initially, the bacterial composition was highly dominated by *Citrobacter amalonaticus, Clostridium butyricum, and Enterococcus xiangfangensis* (Figure 4a). Towards the end of the experiment, the abundance of *C. amalonaticus* and *C. butyricum* decreased and *Ethanoligenens harbinense, Enterococcus olivae, E. xiangfangensis* and *Clostridium pasteurianum* became dominant. Aside from these highly abundant ASVs, 15 other ASVs comprised about a fifth of the total bacterial abundance. These, from now on called rare ASVs, showed independent dynamics from the highly abundant species (Figure 4a and Figure 5). In particular, the abundance of rare ASVs remained stable during the 60 days of the experiment. In contrast, after the heat-shock treatment, only two ASVs were present (*C. guangxiense* and *C. pasteurianum*; Figure 2b), and their abundance remained stable throughout the experiment being *C. guangxiense* the dominant ASV.

**Figure 4.**
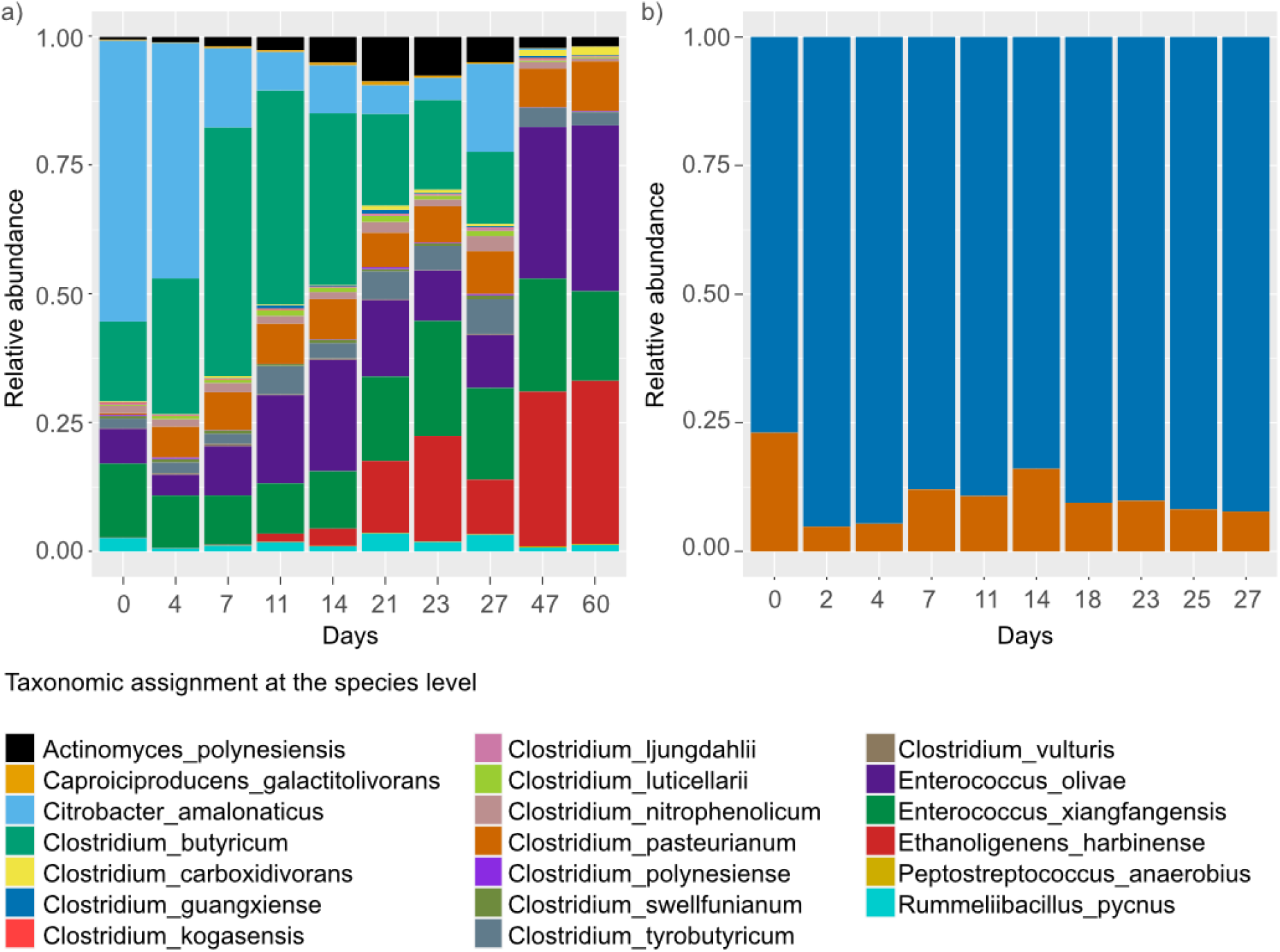
Microbial diversity composition per treatment and over time. Mean relative abundance (N=12) of different amplicon sequence variants (ASVs) in (a) the non-treated and (b) heat-shock treated bioreactors.

**Figure 5.**
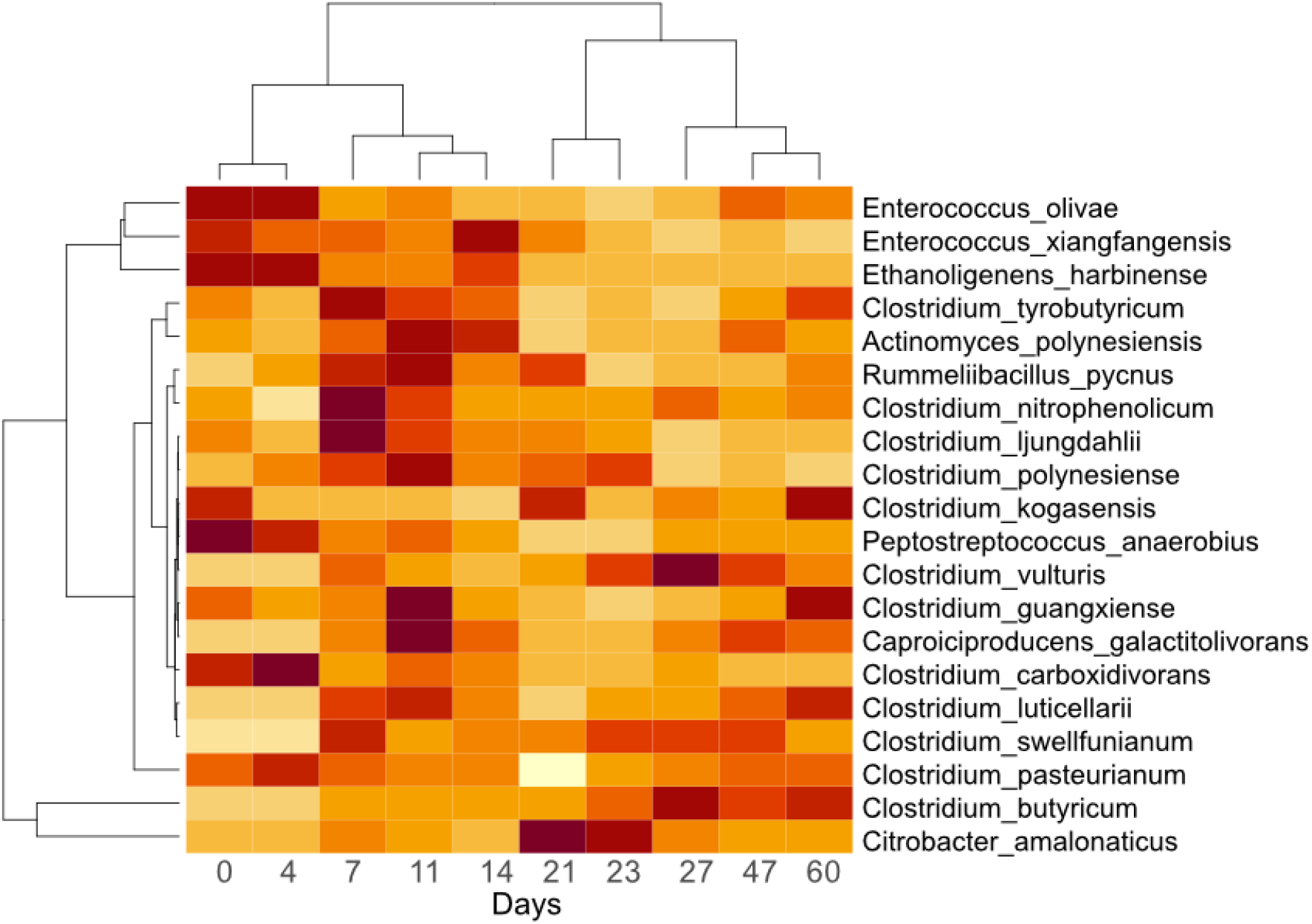
Heatmap showing the variability of the relative bacterial abundance over time in the non-treated bioreactors. Abundance is standardized per ASV to reflect variability between times for each ASV. Each day represents the average abundance per ASV (n = 12).

### Relationship between microbial diversity and function of bioreactors

Since only two species survived in the heat-treated bioreactors and little variation was observed in their function, all correlation and multivariate analyses were performed only in the communities of non-treated bioreactors. It was found that hydrogen production was positively related to the abundance of certain ASVs. For instance, *E. harbinense, C. pasterianum E. olivae, Clostridum carboxidorans* and *Peptostreptococcus anaerobius* are positively correlated with biogas production while *C. amalonaticus, C. butyricum, Clostridium swellfunianum, Clostridium vulturis, Rummelibacillus pycnus* and *Clostridium nitrophenolicum* were negatively correlated (CCA in Figure 6 and Pearson correlations in Figure 7). The bacteria that had negative correlations with biogas production tended to present positive correlations with, either butyric acid and/or propionic acid.

**Figure 6.**
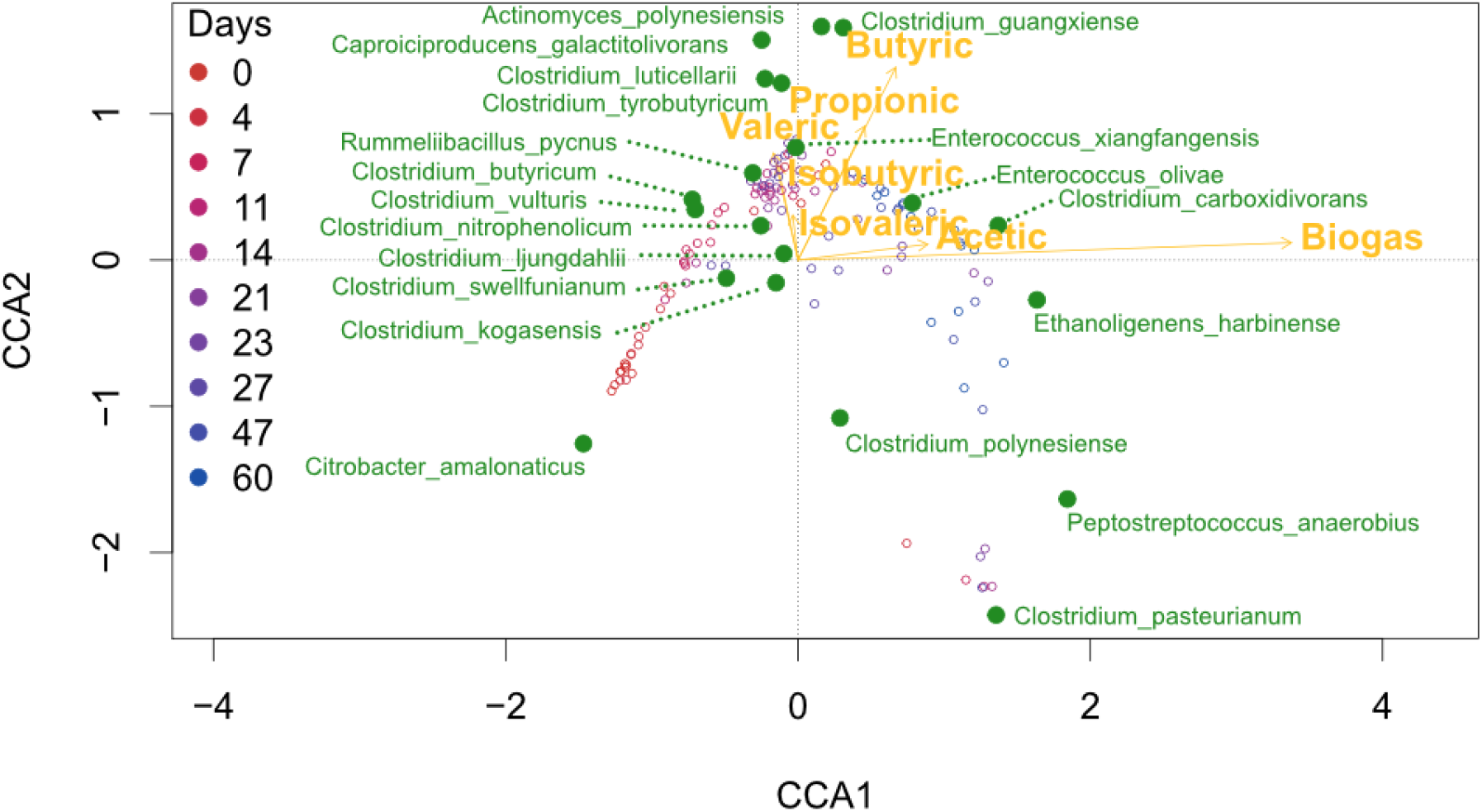
Canonical correspondence analysis (CCA) biplot (n = 120, 12 per day). CCA representing the relationship between bioreactors function (biogas and volatile fatty acids) and microbial composition in the non-treated bioreactors. Yellow points represent ASVs, vectors represent metabolites and hollow points represent samples colored by the time of the experiment when they were taken. Eigenvalues: axis 1 = 0.324, axis 2 = 0.111. Percentage of variance explained: 15.54 (axis 1), 5.33 (axis 2). Cumulative percentage explained: 20.88.

**Figure 7.**
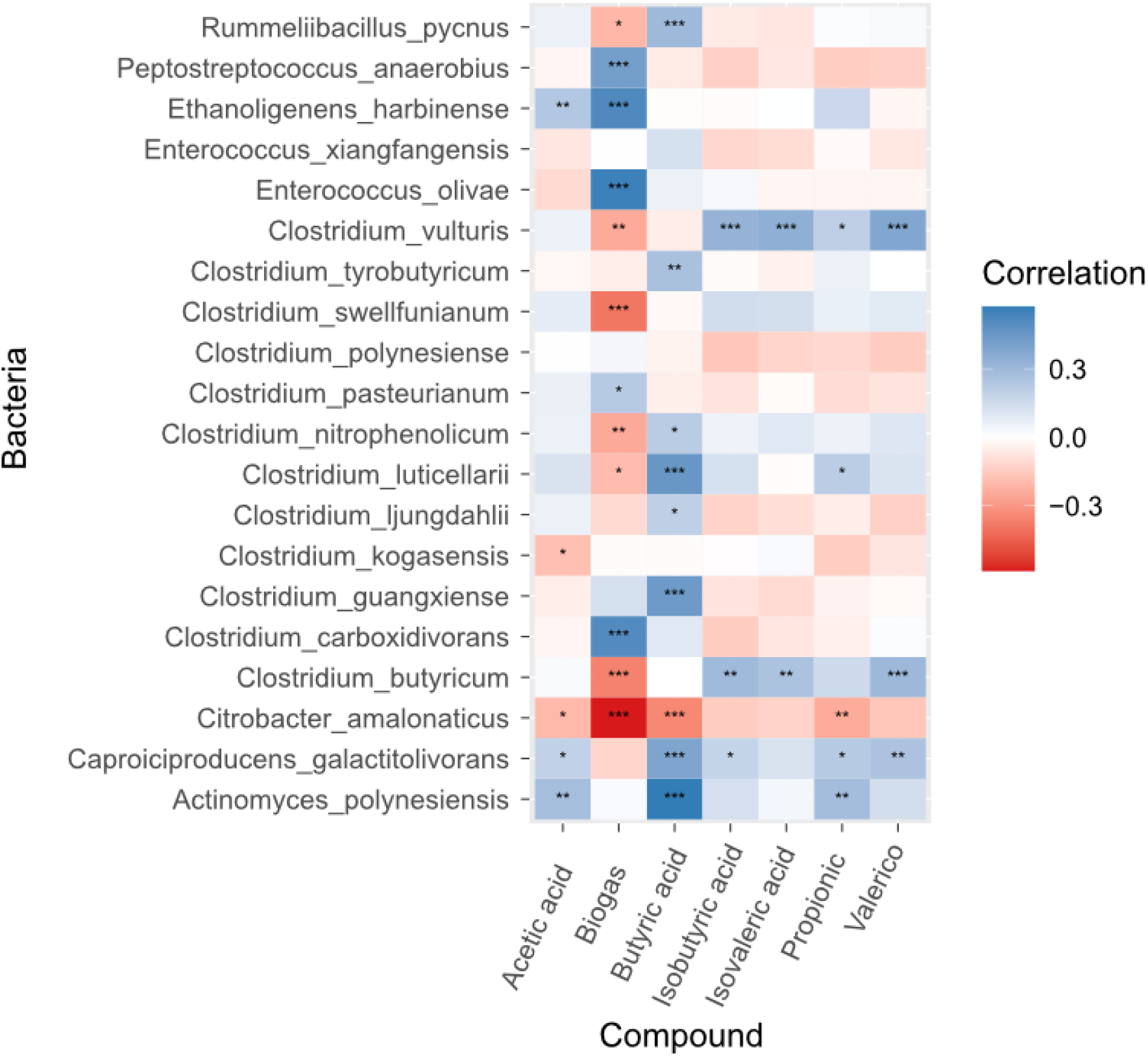
Pearson correlations between microbial abundance, biogas production and volatile fatty acids concentration in the non-treated bioreactors (n= 120). Red shades represent negative correlations and blue shades represent positive correlations. Stars inside each cell represent the significance of the correlation (“***” = 0.001; “**” = 0.01; “*” = 0.05).

For instance, ASVs like *R. pycnus* were positively related to butyric acid, while *Clostridium luticellarii* was positively related to, both, butyric and propionic acids. The rest of the analyzed volatile acids (isobutyric acid, isovaleric acid and valeric acid) and microbial ASVs show no clear relationship to hydrogen production. The CCA (Figure 6), also shows that the structure of the microbial composition of samples in all replicates followed similar composition and function during the experiment. Thus, samples from earlier times in the experiment showed similar microbial structure and functional performance to samples from the end of the experiment.

### Potential ecological interactions

The inferred interactions tended to be potentially beneficial for bacteria that were positively related to biogas (such as *E. harbinense* and *E. olivae*) and potentially negative for bacteria that were negatively related to biogas production (like *C. amanoliticus* and *C. butyricum*; Figure 8). For example, it was inferred that *E. harbinense* and *E. olivae* benefited from *C. pasteurianum*. On the other side, *C. amanoliticus* was negatively affected by *C. pasteurianum* and *R. pycnus, and C. butyricum* was affected by *E. xiangfangensis*. Meanwhile, *C. nitrophenolicum* was a low-abundant bacteria with several interactions that could have modified the consortia composition. For instance, *C. nitrophenolicum* benefited *E. harbinense* but was negatively affected by C. *amanoliticus and C. butyricum*. In turn, *C. nitrophenolicum* benefited from *E. xiangfangensis* and *Clostridium tyrobutyricum*.

**Figure 8.**
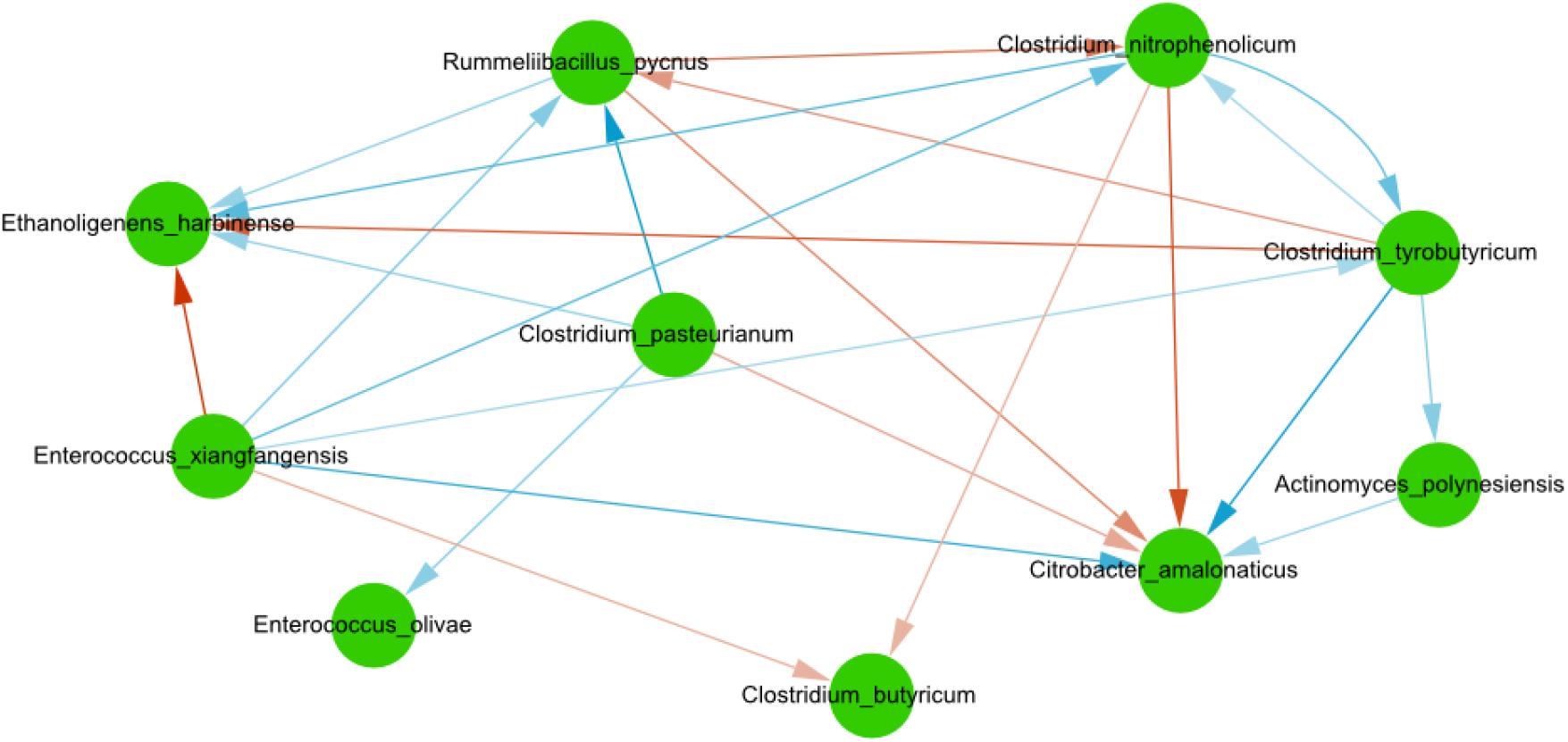
Interactions inferred in the non-treated bioreactors. Nodes denote bacterial species, blue edges denote positive interactions, red edges denote negative interactions, the direction of the arrows represents the direction of the interaction, and the intensity of the color of edges represents the strength of the interaction.

**Figure 9.**
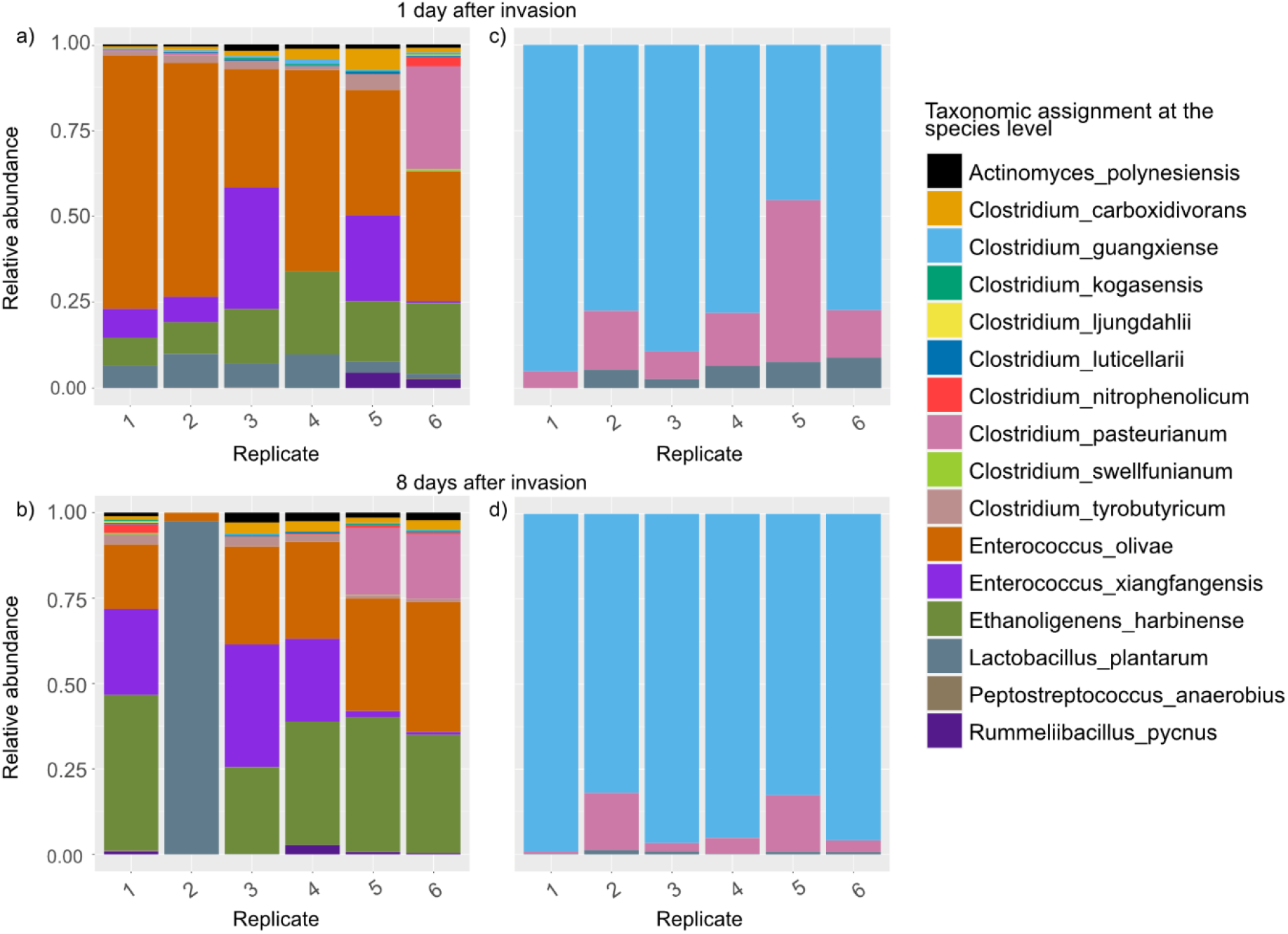
Microbial composition after 6 replicates were divided and inoculated with *L. plantarum*. Panels a and b correspond to the non-treated bioreactors, 1 day and 7 days after the inoculation respectively. Panels c and d correspond to the heat-shock treated diversity bioreactors, 1 day and 7 days after the inoculation respectively.

#### Robustness to invasion

Although both types of consortia were resistant to invasion by *L. plantarum*, each treatment showed different responses to invasion. Figure 7 shows the microbial composition of the bioreactors after the invasion of *L. plantarum* was performed. In the non-treated biroreactors, after 24 hours of the invasion, *L. plantarum* was present in all the bioreactors where it was inoculated. After one week of inoculation, *L. plantarum* was detectable in only one of the bioreactors. Importantly, in the bioreactor where *L. plantarum* was still present, it became the dominant organism causing the collapse of hydrogen production. In the heat-shock treated bioreactors, an initial presence of *L. plantarum* was observed 24 hours after the inoculation. Contrarily to the non-treated bioreactors, *L. plantarum* never became dominant in any of the bioreactors, rather, it persisted in small abundances after one week. At the end of the experiment, all the bioreactors where *L. plantarum* did not establish kept a microbial composition like that observed before the inoculation.

## Discussion

In microbial communities, diversity and function are tightly related although the relationship is not straightforward (Louca et al., 2018; Escalas et al., 2019). Further, microbial composition and function are dependent on, not only the culture conditions but also the biological characteristics of the involved microorganisms. In hydrogen-producing bioreactors, microbial diversity is harnessed to achieve a particular function derived from it. These microbial communities are subjected to a varied set of culture conditions to achieve hydrogen production that have distinctive impacts on their diversity (from mild to drastic microbial diversity reduction by aggressive pretreatments). However, this diversity reduction alters the structure, function and stability of the microbial communities with unknown ecological consequences (Philippot et al., 2013; Wagg et al., 2014; Hernández et al., 2019). In this work, we aimed to analyze the effect of reducing the diversity of microbial consortia on ecological processes (like biotic interactions and robustness).

### Treatments on inoculum predictably affected the diversity, structure and function

To determine how variation in microbial diversity affected function, we periodically tracked the biogas and volatile fatty acid production in both, the non-treated and heat-shock treated bioreactors. Overall, the results show that the bioreactors of both treatments showed distinctive patterns of composition and functional stability (Figures 2 and 3).

Both treatments showed a slow but steady increase in biogas production with a decreased variability at the end of the experiment. In the high diversity treatment, such a pattern was more noticeable. This pattern of ascending biogas production has been observed previously (Kim and Shin, 2008; Kannaiah Goud et al., 2014). The most obvious cause is a shift in microbial composition either to an increased abundance of hydrogen-producing bacteria (Castelló et al., 2018; Palomo-Briones et al., 2018; Yang and Wang, 2019). Also, the higher stability and biogas production in the pretreated treatment are in accordance with the previous observation that pretreated communities have been observed to have better long-term stability by selective enrichment of *Clostridum* species (Singh and Wahid, 2015; Cabrol et al., 2017).

To investigate the microbial composition associated with the observed patterns in the biogas production, we performed the culture-independent molecular characterization of the microbial diversity in ten timepoints in the 12 replicates for each treatment. Figure 4 shows that in the non-treated bioreactors a significant shift of the dominant bacteria occurred. Also, the CCA (Figure 6) showed that there was a shift in microbial composition where a subsample of ASVs (including *E. olivae, E. harbinense* and *C. pasteurianum* and *C. carboxidivorans*) increased in abundance late in the experiment and were positively related with biogas production. The presumed positive relationship of *E. Harbinense, E. olivae, C. pasteurianum* and *C. carboxidivorans* with hydrogen production was confirmed in the correlation analysis (Figure 7). *E. harbinense and C. pasteurianum* are known hydrogen producers (Wang et al., 2009; Masset et al., 2012; Srivastava et al., 2017) and *Enterococcus* species are frequently present in hydrogen-producing consortia and some are hydrogen producers (Pachapur et al., 2015; Braga et al., 2018; Yin and Wang, 2019). *E. olivae* has been reported to produce gas on glucose (Lucena-Padrós et al., 2014) although this gas was not identified so, although possible, it is not clear if *E. olivae* contributed to hydrogen production. On the other side, *C. carboxidivorans* is an homoacetogen capable of growing by consuming H_2_ and CO_2_ (Fuentes et al., 2021). The positive relationship between *C. carboxidivorans* and hydrogen could be the result of its increased growth due to hydrogen availability.

In the heat-treated bioreactors, microbial richness was severely reduced consisting only of *C. guangxiense* (which dominated the bioreactors) and *C. pasteurianum*. Although our case is an extreme example of reduced microbial diversity, other experiments also show very few ASVs (e.g. Muñoz-Páez et al., 2019). It is worth noticing that the stability observed in the heat-shock treated bioreactors of our experiment might be deceiving. When scaling the process to industrial settings, anaerobic conditions and sterility represent important challenges since they are difficult to maintain and not cost-effective (Cabrol et al., 2017) and have been discussed earlier as being the result of selective pretreatments (Hawkes et al., 2002). Our observations tell us that the loss of diversity possibly affected ecological processes in the bioreactors which resulted in different behaviors between treatments. Resolving these ecological processes might help to explain why microbial composition changed in the non-treated bioreactors through time or why in the heat-shock treated bioreactors both ASVs remained with a stable abundance.

### Stability in diversity treatments was the result of distinctive ecological processes

The impact of ecological microbial interactions on composition and function has been studied in several systems. For example, cooperative and competitive interactions have been reported to affect the assemblage of communities in natural and synthetic consortia (Foster and Bell, 2012; Cordero and Datta, 2016). Further, biotic interactions have been shown to influence the function of microbial systems, for example by affecting their growth (Pekkonen et al., 2013), productivity (Fiegna et al., 2015) and resilience (Feng et al., 2017). Also, from an evolutionary point of view, interactions have been reported to evolve in time frames relevant to the operation time of bioreactors (Rosenzweig et al., 1994; Harcombe, 2010; Poltak and Cooper, 2011; Jeffrey Morris et al., 2014). Thus, studying the biotic interactions in microbial communities can help us deduce which organisms had a role in shaping the abundance and function in microbial systems like bioreactors.

In both treatments, we could observe that ecological interactions and derived ecological processes can influence the observed dynamics in contrasting ways. In the non-treated bioreactors, more ASVs meant more chances that interactions with strong effects on composition and function occurred for a longer time. For instance, in the non-treated bioreactors, negative (e.g. *C. pasteurianum* towards *E. harbinense* and *E. olivae*) and positive interactions (e.g. *Clostridium tyrobutyricum* and *Enterococcus xiangfangensis* towards directed towards *E. Harbinense*) may be responsible for the increase of abundance *E. harbinense* and *E. olivae* in the last part of the experiment. Interestingly, *Enterococcus* ASVs frequently coexist with *Clostridium* and *Ethanoligenens* bacteria in various environments that range from human hosts to bioreactors (Wang et al., 2009; Pachapur et al., 2015; Valdez-Vazquez et al., 2015; Braga et al., 2018). As such, our data indicates that the variable roles reported for lactic acid bacteria (like *Enterococcus*) in hydrogen-producing consortia (Sikora et al., 2013) are ASVs-specific. For example, in our experiment other enterococci (*E. xiangfangensis*) had a negative relationship with, both, *E. harbinense* and *C. butyricum*. Lastly, low-abundant bacteria (like *C. nitrophenolicum, Rummeliibacillus pycnus* and *Clostridium tyrobutyricum*) were involved in interactions that could favor the change in the overall composition. Low-abundance bacteria have been observed to be able to affect the whole structure of microbial communities either by acting as keystone species, being highly active by acting as reservoirs against environmental perturbations and should not be overlooked (Rafrafi et al., 2013; Shade et al., 2014; Lynch and Neufeld, 2015).

Contrarily, in the case of the heat-shock treated bioreactors fewer ASVs led the consortia to reach stability more quickly. The compositional and functional stability was presumably the result of the two ASVs showing a mostly neutral interaction. We deduce this neutrality since the two ASVs coexisted without many changes in abundance during the whole experiment. Also, based on the interactions inferred for the non-treated bioreactors, no significant interaction was found between *C. pasteurianum* and *C. guangxiense*. This agrees with observations in other communities, both natural and artificial, where several species of *Clostridium* can coexist (Graf et al., 2015; Thi Hoang et al., 2018). Since not every *Clostridium* (or for that matter any bacterial group) contributes equally to function, our results draw attention to fully understanding the reasons behind the presence of certain species in microbial consortia. In the case of the heat-shock treated bioreactors, the presence of *C. pasteurianum* and *C. guangxiense* might be the result of deterministic processes like different spore germination times and growth rates that have been noted to be important in community assembly (Hawkes et al., 2002; Elke Jaspers and Jorg Overmann, 2014). Also, stochastic processes like drift could have been important as has been noted earlier for the assembly of microbial communities in bioreactors (Liébana et al., 2019). As mentioned before, the drastic reduction in ASVs in the heat-shock treated bioreactors might give the impression of increased stability in the bioreactors. However, in addition to the fact that strict anaerobic conditions had to be used, the diminished diversity can lead to less long-term instability, due to, for example, a lack of functional redundancy in case of perturbations (Louca et al., 2018).

Invasion by competitive species in non-sterilized substrates is one of the reasons for bioreactor’s instability (Castelló et al., 2018). Several factors have been shown to influence the resistance to invasions that microbial communities. For example, diversity has been positively linked with a reduced probability of invasion while stochastic processes have been linked to invasion success (Mallon et al., 2015). In this line, the function and structure of the microbial communities of the two treatments showed different responses to invasion (the effect on biogas production and the ratio and success of invasion) that could be related to ecological processes specific to each of them. In the non-treated bioreactors, only in one replicate, the invader became dominant while in the rest of the replicates no presence of the invader was observed after one week. Contrastingly, in the heat-shock treated bioreactors, *L. plantarum* established in at least 4 of them (although in low abundances) but never dominated. Similarly, biogas production was affected more severely in the heat-treated bioreactors, indicating that the alien species had a competitive effect on the microbial communities of such bioreactors. As, such, in the heat-shock treated bioreactors *L. plantarum* might have managed to exploit an available niche which is an important factor regarding establishment of invaders (Kinnunen et al., 2016). In the non-treated bioreactors, since the establishment of *L. plantarum* occurred only once, it is possible that random processes like drift contributed to invasion success in the particular bioreactor where it established (Kinnunen et al., 2016). Additionally, the high effect of *L. plantarum* in the community might be explained by similarity in fitness with the resident community which might have increased the invasion effect (Li et al., 2019). Thus, our results show that the relationship between invasion success and microbial diversity is complex even in simple communities. Further studies must be performed using complex substrates (like agro-industrial effluents) and varying conditions like those intended to be used in industrial-scale bioreactors.

## Conclusions

Despite a long line of evidence showing that diversity has a strong influence on the function and stability of bacterial communities, it is not clear how ecological mechanisms (from which function and stability are dependent) are related to diversity and how they are affected when diversity changes. The results in this work show that the establishment and medium-term behavior of bacterial communities are the outcomes of the interplay between the number and identity of ASVs, biotic interactions and culture conditions. In particular, we found that: (1) higher diversity slowed the stabilization of microbial abundance and dynamics but was ultimately predictable as biotic interactions were important for the assembly of microbial consortia, (2) loss of diversity made stabilization faster but less resilient against adverse culture conditions (aerobic conditions) and (3) higher diversity made invasion success less probable. Further studies investigating the molecular basis of interactions and confirmatory experiments on the nature of the interactions and their effect could help identify mechanisms and processes that caused changes in function and invasion behaviors.

Finally, due to the importance of controlling diversity in biotechnological settings, this study provides a baseline to incorporate ecological theory and experimental designs into the study of biotechnological settings like bioreactors. Further, this type of study might increase the knowledge of the relationship between function and diversity which in turn will help improve the yield and stability of hydrogen-producing bioreactors.

## Acknowledgements

We thank Marisol Perez Rangel for technical assistance during bioreactors establishment and for providing the strain *Lactobacillus plantarum* used in this work. The authors acknowledge Adalberto Noyola Robles and Margarita Cisneros Ortiz for her support in the laboratory work at the Laboratorio de Ingeniería Ambiental, Instituto de Ingeniería, UNAM. Marcelo Navarro-Díaz is a doctoral student from the Programa de Doctorado en Ciencias Biomédicas, Universidad Nacional Autónoma de México (UNAM), and was supported by Consejo Nacional de Ciencia y Tecnología (CONACyT), scholarship no. 594615. Financial support of this work was received from the “Fondo de Sustentabilidad Energética SENER-CONACYT (Mexico)”, Bioalcohols Cluster (Grant No. 249564).

